# G-IRAE: a Generalised approach for linking the total Impact of invasion to species’ Range, Abundance and per-unit Effects

**DOI:** 10.1101/2021.10.18.464772

**Authors:** Guillaume Latombe, Jane A. Catford, Franz Essl, Bernd Lenzner, David M. Richardson, John R. U. Wilson, Melodie A. McGeoch

## Abstract

The total impact of an alien species was conceptualised as the product of its range size, local abundance and per-unit effect in a seminal paper by Parker and colleagues in 1999, but a practical approach for estimating the three components has been lacking. Here, we generalise the impact formula and, through use of regression models, estimate the relationship between the three components of impact, an approach we term G-IRAE (Generalised Impact – Range size – Abundance – per-unit Effect). Moreover, we show that G-IRAE can also be applied to damage and management costs. We propose two methods for applying G-IRAE. The species-specific method computes the relationship for a given species across multiple invaded sites or regions, assuming a constant per-unit effect across the invaded area. The multi-species method combines data from multiple species across multiple sites or regions to calculate a per-unit effect for each species. While the species-specific method is more accurate, it requires a large amount of data for each species. The multi-species method is more easily applicable and data-parsimonious. We illustrate the multi-species method using data about money spent managing plant invasions in different biomes of South Africa. We found clear differences between species in terms of money spent per unit area invaded, with per-unit expenditures varying substantially between biomes for some species. G-IRAE offers a versatile and practical method which can be applied to many different types of data, to better understand and manage invasions.

## 1. Introduction

Many invasive alien species cause deleterious environmental and socio-economic impacts and are among the main threats to biodiversity (Bellard et al. 2016; Maxwell et al. 2016; Seidl et al. 2018; IPBES 2019). Biological invasions cost the global economy at least a thousand billion US dollars each year – as a result of direct economic damage and money spent on damage mitigation (e.g. invader control, ecosystem restoration) (Diagne et al. 2020, 2021). Several frameworks have been developed and applied globally to better conceptualise and characterise different types of impacts and costs (e.g. Blackburn et al., 2014; Jeschke et al., 2014; Kumschick et al., 2020; Van der Colff et al., 2021). Despite these advances, further work is needed to disentangle the different components of impacts and enable their objective quantification.

To better understand impacts, Parker et al. (1999) proposed a formula in which the total impact of an alien species is calculated as the product of its spatial extent (range size), local abundance, and per-unit effect (the impact caused by a single individual, unit of biomass or unit of invaded area). The positive relationship between total impact and range size or local abundance is straightforward; the more individuals or the greater the invaded area, the greater the impacts. These relationships are often non-linear (Bradley et al. 2019), and these variables can increase at different rates after species introduction (McGeoch and Latombe 2016; Latombe et al. 2020; Cheney et al. 2021). Per-unit effects are less well understood. Most attempts to disentangle the three components of alien species impact and account for non-linearities in these relationships to date have been theoretical and conceptual (e.g. Yokomizo et al. 2009; Thiele et al. 2010; Buckley and Catford 2016; Vander Zanden et al. 2017). These studies generally assume that the per-unit effect changes non-linearly with species range size and abundance, and discuss the specificities of these relationships and which factors may influence them (e.g. Strayer, 2020). Density-impact relationships have sometimes been assessed empirically, but this has been done for specific taxa in different locations independently, using a variety of approaches. The comparison between these relationships across multiple taxa, although insightful, remains qualitative due to the lack of a common quantitative approach (Norbury et al. 2015).

Parker et al.’s (1999) publication has been highly cited (2124 citations on Google Scholar as of 2021-09-30). However, despite it being one of the most highly cited papers in invasion science (Pyšek et al. 2006), and despite the formula’s potential utility for comparing, ranking, and prioritising species, practical applications of the concepts that underpin Parker et al.’s (1999) formula are still missing more than 20 years after it was proposed (cf. the uptake of other concepts in invasion science, Wilson et al., 2020). Here, we propose the G-IRAE (for Generalised Impact – Range size – Abundance – per-unit Effect) approach that combines Parker et al.’s (1999) formula and more recent perspectives on the relationship between density and impact or cost to effectively compare the components of impacts of multiple alien species using real data. We expand and linearise Parker et al.’s (1999) formula, enabling us to use linear models (and associated statistical tools) to calculate the independent components of impact (i.e. a constant per-unit effect of alien species and parameters describing the form of the relationship between impact, range size and local abundance). Better understanding of the differences in per-unit effects among species and of relationships between impact, range size, and abundance is needed to improve management decisions. Quantifying these components will also pave the way to better understand how different attributes of alien species and the environment interact to contribute to the different facets of impact.

Parker et al. (1999) formula focuses on ecological impact. However, we argue (and illustrate in the following) that the concepts can be readily extended to encompass damage costs or management costs, e.g., by considering per-unit management cost or expenditures rather than per-unit ecological impact. The approach can be applied using different indicators (or metrics) of cost and impact, as long as the relationship with range size (or managed range size for management costs and expenditures) and local abundance monotonically increases, to compute per-unit effects, costs and expenditures. As such, G-IRAE provides a method to highlight systematic differences in the amount spent controlling different types of invasions, even when total management expenditures are established a priori due to practical constraints. When applied to the money spent on management, the purpose of these analyses is not, of course, to de facto determine the efficacy of management, but they do provide an important tool to assist conservation practitioners, managers and decision-makers in understanding how the allocation of resources are implemented in practice.

We propose two different methods to implement G-IRAE — the species-specific method and the multi-species method. The species-specific method estimates the per-unit effect and parameters for range size and local abundance for a given species based on data on range size and local abundance for different sites or regions. It is in line with existing approaches based on the density-impact relationship (e.g. Norbury et al., 2015; Strayer, 2020; Thiele et al., 2010; Yokomizo et al., 2009), and allows for a detailed comparison between species. Although useful for capturing and conceptualising differences in the impacts of different alien species, the applicability of this method is limited by data availability. By contrast, the multi-species method can be applied in the context of simultaneous invasions by multiple alien species across multiple sites or regions, with each species characterised by a single combination of total impact or cost, (treated) range size, and average abundance in each invaded or managed site or region. This method assumes a similar relationship with range size and local abundance over the set of alien species under consideration; it is more feasible in practice as the required data are more readily available.

In this paper, we use a case study of alien plant invasions in South Africa to illustrate the multi-species method. Specifically, we combine data on the money spent controlling 18 alien plant species in nine biomes with data on the distribution and abundance of these species, in order to examine how per-unit expenditures on management differ between species in different biomes, based on how resources are allocated between target taxa. We discuss how the general approach introduced here is applicable to (arguably more complex) measures of socio-economic and ecological impacts, and suggest ways of overcoming limitations of the proposed methods, including data collection.

## 2. Methods

### 2.1. Theoretical concepts of per-unit effects formula: the G-IRAE approach

Parker et al. (1999) introduced the following formula to relate the total impact I of an alien species to its range size R, abundance per-unit area A across range R (referred to as local abundance hereafter) and the per-unit effect E (i.e. per-unit abundance and per-unit area):

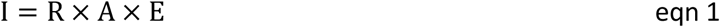

The units of range size and abundance, and the type of impacts can vary. Range size can be expressed as ha or m^2^ or occupancy of particular sites; units of abundance can be numbers of individuals, biomass, or percent coverage; and units of impact might be financial or in terms of some index of environmental damage caused (e.g. biodiversity intactness or the environmental impact classification for alien taxa; see Discussion for more details). If I represents the total money spent on management, R the treated range size, and A the local abundance across the treated range, then E is the management expenditures per-unit abundance and per-unit area treated (and variation in E across sites or taxa will be of interest to those monitoring and planning management).

Equation 1 assumes that I increases linearly with R and A. However, depending on how impact is characterised, the relationships is more likely to be non-linear. For example, the relationship between the abundance of alien species and native species diversity is often negative and convex (Bradley et al. 2019), i.e. there is a non-linear relationship between the abundance of alien species and the absolute loss of native species diversity, which is one possible measure of impact. This has often been interpreted as a variation in per-unit effect with R and A. That is, equation 1 can be reformulated as:

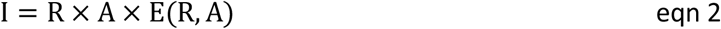

Four typical, theoretical relationships between I, E, R and A have been proposed to develop the concepts behind equations 1 and 2 (Yokomizo et al. 2009; Thiele et al. 2010; Buckley and Catford 2016; Vander Zanden et al. 2017) (Fig. 1). Type I curves assume a linear relationship between Iand Ror A, and therefore a constant per-unit effect E, independent of R and A. Type II curves assume that the per-unit effect E of a species is high at low range size and abundance values, but decreases as range size and abundance increase. Type II curves therefore lead to a rapidly increasing Impact at low R and A values, which tends to saturate at high values. Type III curves assume the opposite of type II curves that is that per-unit effects are low at low range size and abundance, but increase as range size and abundance increase. Under type III relationships, alien species must reach a certain range size or abundance before substantial (measurable) impact manifests. Finally, type IV curves are a mix of type II and III curves, with a slow increase in impact at low R and A values, which temporarily accelerates before tending to saturate at high R and A values.

**Figure 1.**
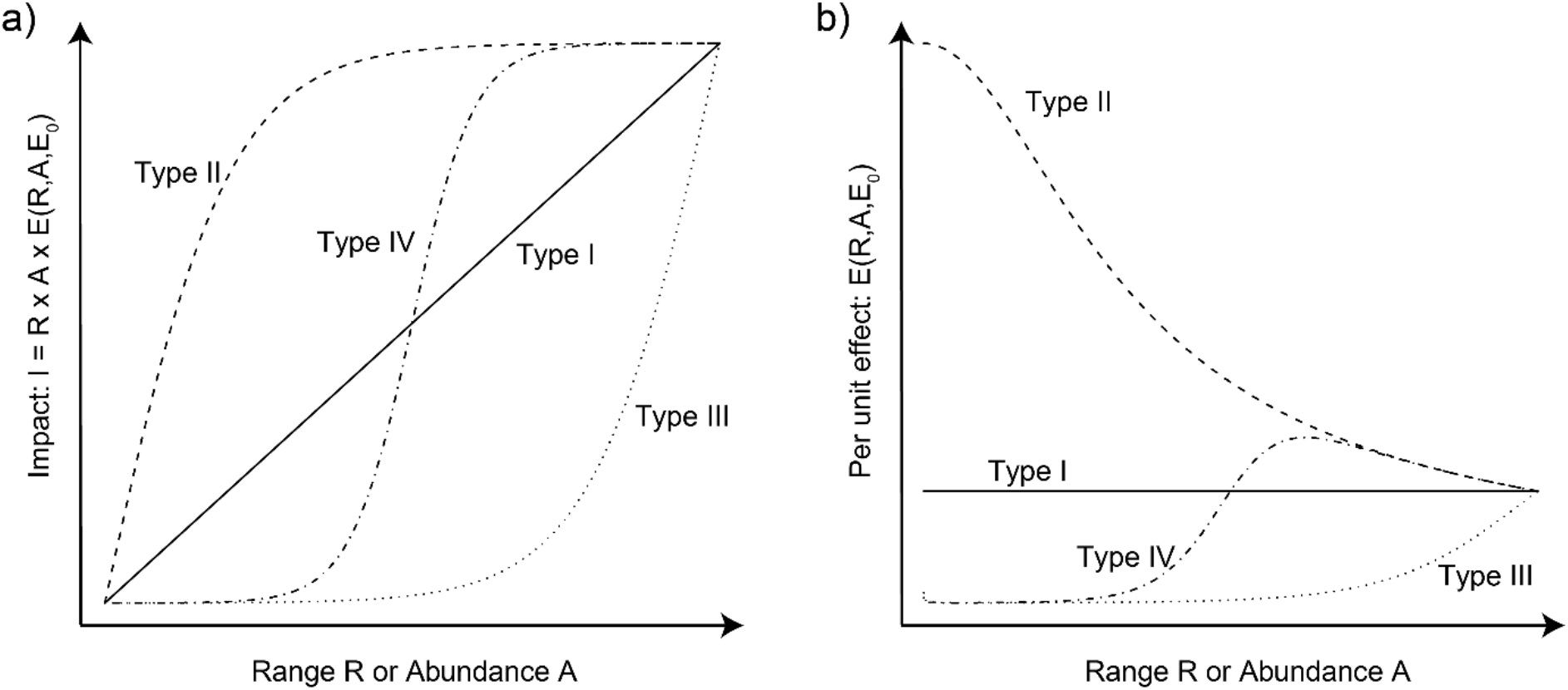
Four archetypes of relationships between total Impact (I), Range size (R), Abundance (A) and per unit effect E of an alien species following the formula proposed by Parker et al. (1999). a) The relationship between I, R and A is often non-linear. Note that a different curve can describe the relationship between Iand R and between Iand A, i.e. I can increase differently with R and A. Ican then be computed as the product of the values for a given (R, A) combination. b) The non-linearity is generated by the relationship between the per-unit effect E and R, A and the constant E_0_ (as for I, the relationship can differ for R and A). The shapes of the relationship are for illustrative purposes only; the exact shapes will vary depending on the system and the method used to estimate them.

These relationships are useful for conceptualising how impacts and per-unit effects scale with changes in range size and local abundance of alien species. Although they have been modelled in different studies using a variety of approaches and combinations of variables (Norbury et al. 2015), a general approach to apply them quantitatively across species and environments has yet to be proposed (but see Cuthbert et al. (2019) for the related concept of functional response). To do so, we propose to generalise equation 1 as follows:

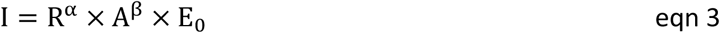

Where E_0_ is the per-unit effect of a species for one unit of range size and abundance; it is therefore constant for a given species, contrary to the interpretation of equation 2. Note that equation 3 can also be written as follows, to account for the fact that per-unit effects can vary with range size and local abundance, as conceptualised in equation 2:

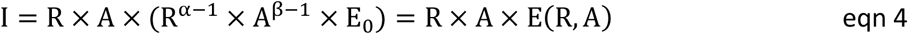

The α and β exponents enable us to capture some non-linearities between total impact, range size and local abundance. In particular, we can model three of the four theoretical relationships described above (Fig. 1). Unit values for α and β indicate a linear relationship between impact, range size and abundance, i.e. a type I relationship. Values of α and β below one but greater than zero lead to a deceleration in the increase of impact with range size and local abundance, and therefore corresponds to a type II relationship, noting that this functional form does not allow impact to truly saturate, but only to decelerate as R and A increase. Values of α and β greater than one leads to an acceleration in the increase of impact with range size and local abundance, and therefore corresponds to a type III relationship. Values below 0 are not considered here because, α = β = 0 would indicate a constant impact, regardless of range size and abundance, and α, β < 0 would indicate that impact decreases with range size and abundance, which is unrealistic.

Using a log-transformation, we can linearise equation 3 as follows:

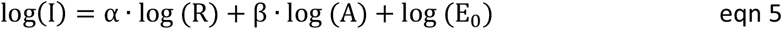

Linearising equation 3 enables us to use powerful statistical tools, including linear models and mixed effects linear models, to estimate the parameters α and *β*, and the per-unit effect constant E_0_, while accounting for statistical significance, as we develop below.

Although type IV relationships can be modelled mathematically by a sigmoid function as 1/(1 + exp (−α · (R− R_0_)), the product of sigmoid functions would not allow for a linearisation after a log-transformation, as in equation 5, and so are not considered further here.

### 2.2. Species-specific method

Ideally, equation 5 would be applied to species individually, to obtain a unique relationship for each species. Applying equation 5 to a single species would require data on impact, range size and local abundance in different locations where the per-unit effect is assumed to be driven by the same factors, and therefore to have the same relationship with R and A. In that case, log(E_0_) for a given species is simply the intercept of the following linear model:

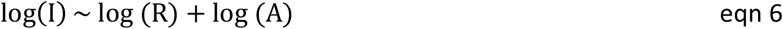

Applying equation 6 to individual species separately would generate comparable α, β, and E_0_ values between species, allowing us to predict when a species will have a greater or lower impact than another, and therefore to better prioritise management actions (see Fig. 2 and Discussion). This approach could therefore be applied to datasets with comprehensive and comparable data on species range, abundance and impact in multiple locations, but we know of no such datasets for multiple species.

**Figure 2.**
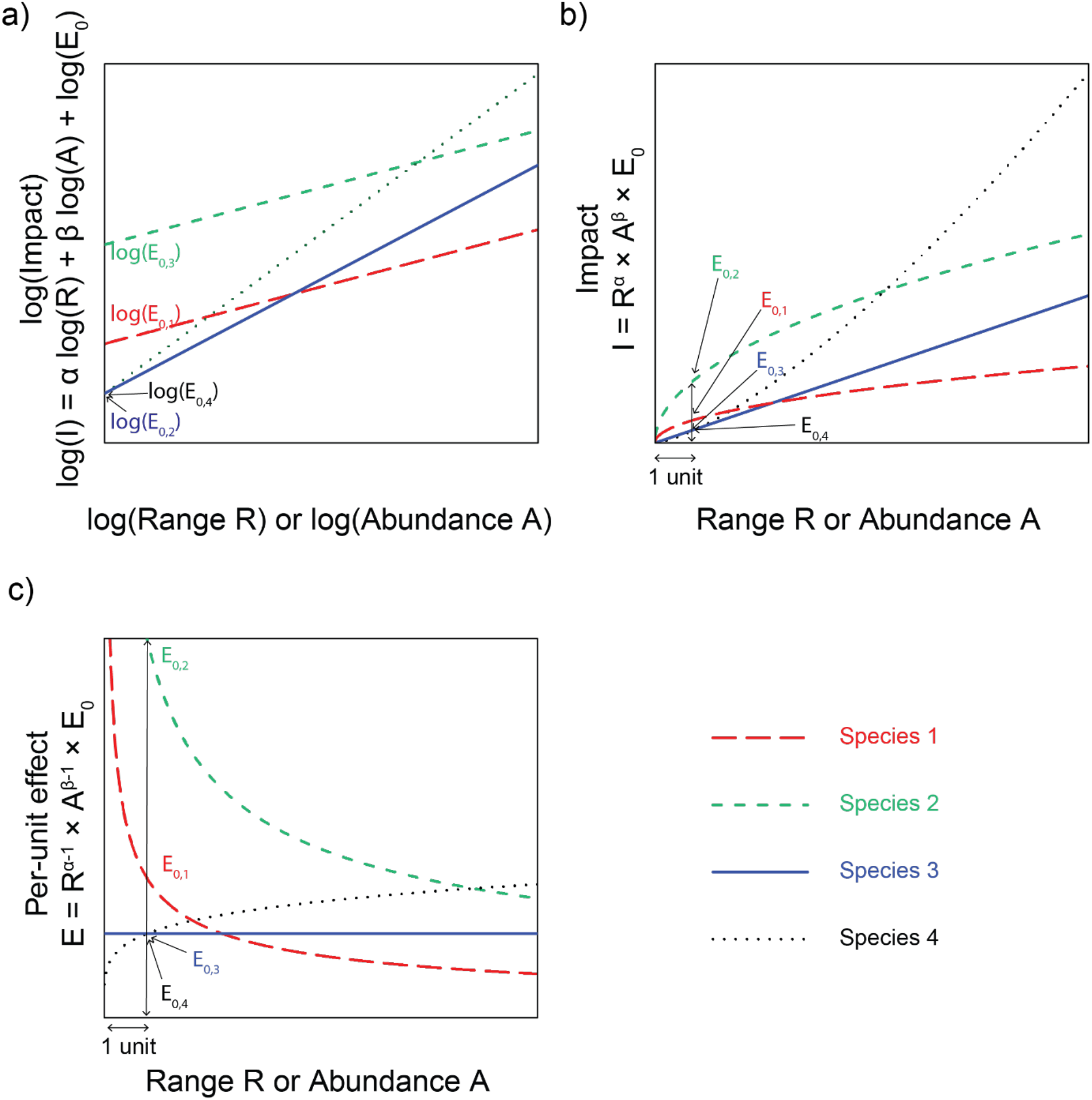
Theoretical illustration of the calculation of the relationship between impact I, per-unit effect E, range size R and local abundance A of alien species, following the species-specific method that computes the relationship for each species independently. a) For each species, the relationship between log(I), log(R), log(A) and log(E_0_) is computed using a linear model. The α and β coefficients can take different values: between 0 and 1 (species 1 and 2; type II); equal to 1 (species 3; type I); or above 1 (species 4; type III). For each species, log(E_0_) is the intercept. Note that the values of α and β will likely be different for the same species, and a species will be characterised by a combination of two slopes, i.e. a surface, simplified here as a one-dimensional slope for clarity. b) Exponential-transforming the relationship reveals that, for this theoretical example, the relationship decelerates for species 1 and 2, and that the lower E_0_ value for species 1 leads to a quicker deceleration. The unit value for the coefficients of species 3 leads to a linear relationship, whereas the relationship accelerates for species 4. For each species, E_0_ is the value of Ifor one unit of R and A. c) These relationships can be expressed as the relationship between the per-unit effect of each species and their range size or abundance. The per-unit effect of species 1 and 2 decreases with range size and abundance, whereas it increases for species 4, and is constant for species 3.

### 2.3. Multi-species method

Multiple alien species often co-occur in a given area, but can have different spatial distributions within this area, and different impacts (McGeoch and Latombe 2016; Cheney et al. 2021). When each species is characterised by a single I, R and A value in a given area, these data can be combined across species to calculate a measure of per-unit effect per species. Given a set of multiple species *s*, and if we assume that the relationship between the total impact I, R and A is the same for all species included in the analyses (but see Discussion for further details on this topic), equation 5 can be reformulated as follows:

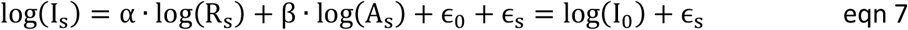

Where ϵ_s_ is the residual for species s, after computing log(I_0_), the estimation of log(I_S_) assuming the same per-unit effect ϵ_0_ for all species (i.e. the baseline impact). In other words, ϵ_0_ is the intercept of the linear model:

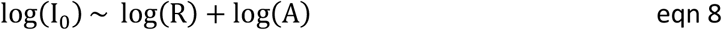

The logarithm of the per-unit effect of each species s, log(E_0,s_), therefore corresponds to the sum of the intercept ϵ_0_ plus the residuals ϵ_s_ after fitting equation 8, as shown in equation 9 (Fig. 3).

**Figure 3.**
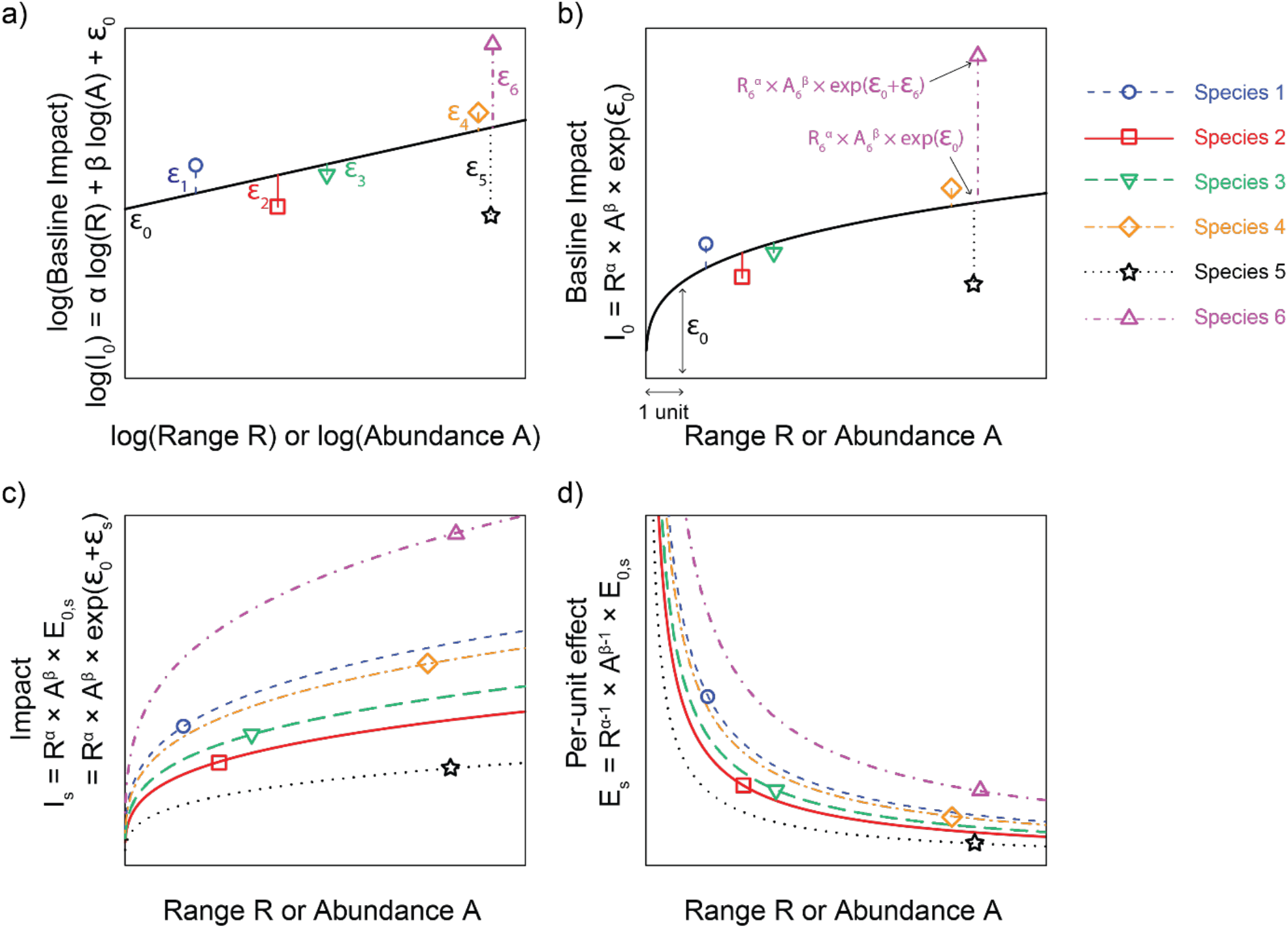
Theoretical illustration of the calculation of the relationship between impact, per-unit effect, range size and abundance of alien species, following the multi-species method that computes the relationship using data from all species simultaneously, when each species is characterised by a single (I_s_, R_s_, A_s_) combination, contrary to the species-specific method (Fig. 2). a) The relationship between log(I_0_), log(R), log(A) and log(E_0_) is computed using a linear model based on the combined data from all species. As a result, only one combination of α and β coefficient values, and a single intercept ϵ_0_, are computed for all species. Note that a combination of α and β values would general a two-dimensional surface, simplified here as a one-dimensional slope for clarity (with a slope in [0,1[for the sake of the example, but it can be above 1). The resulting I_0_ variable represents the baseline impact across species. The difference between the logarithm of the impact of each species log(I_s_) and log(I_0_) is indicated by the residual values ϵ_s_. b) The relationship can be exponential-transformed to reveal how the observed impact values I_s_ differ from the baseline I_0_. Since the α and β coefficient values are below 1 in this example, the baseline I_0_ decelerates as R and A increase. c) A relationship is computed for each species, which only differ from each other in the E_0,s_ values. Since all species share the same α and β values, the I_s_ values decelerate as R and A increase for all species. d) These relationships can be expressed as the relationship between the per-unit effect of each species and their range size or abundance, which decreases for all species.

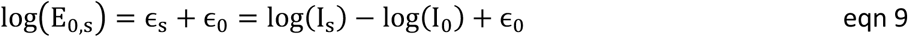

### 2.4. Data

#### 2.4.1. Management costs as a across sites and taxa

Equation 1 was developed with ecological impacts in mind, but is also appropriate for considering other metrics, e.g. economic impacts or management costs (Parker et al. 1999). Our generalisation (equation 3) can therefore encompass multiple types of effect, including the economic impacts of invasions and the amount spent on their management. The amount spent managing a population of an alien species can indeed be considered to depend on the cost of managing one individual or unit of biomass, and on the range size and local abundance treated. The larger the area to manage (irrespective of whether this area is determined a priori or depends on the total area occupied by the invader), the costlier it is likely to be to pursue management, as management actions involve moving people and equipment. Similarly, the more abundant a species is at a given location, the costlier it is likely to be to remove, because of the increase in time required (although this relationship will depend on the management approach used). These relationships can then accelerate or decelerate (i.e. increasing or decreasing per-unit management cost, equation 4, Figure 1), depending on the logistics and approaches used to manage the alien species, the existence of fixed costs (e.g. start-up costs), etc.

The amount of money spent on the management of a particular species is nonetheless often decided a priori, and is influenced by many factors (Panetta 2009). Note that determining a priori the level of funding dedicated to the management of a particular species or the range over which it should be managed has no impact on the approach. Assuming the relationship between I, R, A and E_0_ can be correctly assessed from equation 6 and equation 8, data on I, R and A will enable to estimate E_0_ regardless of their value. For example, if a small overall amount of money I is spend over a large area, E_0_ will be small, indicating that management will likely be inefficient at removing individuals. Reciprocally, if management is designed so that each treated individual is effectively removed, but little money is dedicated to overall management of a species, the treated area will be narrow. In other words, it is possible to see variation in how money is spent even if such money is not spent effectively and the invasion may not actually be managed.

To explore the available data on management costs over South Africa, we evaluated InvaCOST, the most comprehensive database on economic costs of alien species worldwide, which considers both damage and management costs (Diagne et al. 2020). InvaCOST lists 31 studies in South Africa, which provided a comprehensive overview of the economic data available. Amongst these studies, due to issues of comparability, only van Wilgen et al. (2012) provides comparable estimates of the amount of money spent managing 18 taxa (species, genera or families) in nine biomes in South Africa by the Working for Water program between 1995 and 2008, in South African Rands. We therefore use these data to demonstrate the approach following equations 7-9.

#### 2.4.2. Species distribution

Fitting equation 8 requires data at fine scale on range size and abundance of alien species over a large region. van Wilgen et al. (2012) reports the estimated occupied area by these species per biome in 1996 and 2008, obtained from three independent studies (Le Maitre et al. 2000; Kotzé et al. 2010; Van den Berg et al. 2013), and the area treated by the Working for Water program between 2002 and 2008 (all expressed in condensed area, i.e. the treated area effectively occupied by the species, rather than the total extent of occurrence over which the species was managed; see Discussion for details about different measures of range size). We used the area treated as a measure of range size in equation 8.

For abundance data, we used estimates from the Southern African Plant Invaders Atlas (SAPIA; Henderson, 2007; Henderson and Wilson, 2017). SAPIA is the most comprehensive database on alien species distribution in South Africa. It comprises records gathered by 670 participants since 1994, in addition to road surveys by the lead author since 1979 over the whole country (we only kept surveys prior to 2008, which is the latest year of the Working for Water assessment reported in van Wilgen et al., 2012). SAPIA records are, with the exception of the earliest records, geo-referenced to point localities, but notes were taken as to the abundance of invasive plants at the landscape scale (and so at resolutions in the range 1–10km). Abundance was scored as present: abundance unknown or not recorded; rare: one sighting of one or a few plants; occasional: a few sightings of one or a few plants; frequent: many sightings of single plants or small groups; abundant: many clumps or stands; and very abundant: extensive stands. Records whose abundance was scored as present were removed from the analyses since it can belong to any of the other categories. We converted the remaining five classes to an exponential scale (exp(1) to exp(5)) assuming that increases in abundance would be similar to those described in Wilson et al. (2018) for four categories (<2%, 2-10%, 10-50%, >50%). Note that these values (exp(x)) need not represent the actual number of individuals or biomass units; only their relative values are required, as discrepancies with actual values will be accounted for by parameter β in equation 5 (see the Discussion for further details on this topic). For each record, we extracted the biomes in which it was recorded based on its coordinates and the 2006 vegetation map of South Africa, Lesotho, and eSwatini (previously Swaziland) (South African National Biodiversity Institute, 2006), as 2006 is encompassed in the period over which alien species were managed and should therefore best correspond to the biomes reported by van Wilgen et al. (2012).

Many factors come into play when selecting invaded areas to be treated, including the level of invasion, accessibility, native biota, etc. (Roura-Pascual et al. 2009, 2010). We lacked information on how management sites were selected in our case study. Therefore, instead of using the average abundance over all records as our measure of abundance in equation 5, we sampled records for each species and biome, using three different sampling approaches, to assess the sensitivity of the results to site selection. We sampled records based on the proportion Parea_S,B_ of treated area over the total area occupied by the species over the 1996 – 2008 period. The total occupied area was calculated as the maximum of the area occupied in 1996 (i.e. before management) or the area occupied in 2008 plus the area treated between 2002 and 2008 (i.e. the area after management plus the treated area), as reported by van Wilgen et al. (2012) (as natural growth of alien species still occurs during treatment, area in 2008 could be more important than area in 1996 despite treatment, in addition to the fact that areas were estimated differently in the two studies). Under the first sampling approach, we assumed management effort was R andomly allocated across invaded sites, so for each species and biome, Parea_S,B_% of the records were sampled R andomly (restricting the selection to the Northern Cape province for *Prosopis* species, as indicated in van Wilgen et al. (2012)), and the average abundance of the records was reported. Under the second approach, we assumed that management prioritised areas that were densely invaded, as alien species should have the highest ecological impact in such sites (Shackleton et al. 2017). We used observed abundance of records (exp(1) to exp(5)) as a weight when sampling, and reported the average abundance over the records selected using this weighted sampling. Under the third approach, we assumed that the least invaded areas (i.e. those with sparse stands of invaders over large areas) were more likely to be treated, as such sites would be more easily managed and more likely to limit invasive spread, leading to greater management efficacy. We used the inverse of the observed abundance of records (exp(−5) to exp(−1)) as a weight when sampling, and reported average abundance. We only used these three sampling approaches to assess the sensitivity of the results, due to the lack of information on site selection, but other criteria could be applied (Shackleton et al. 2017).

### 2.5. Calculation of the per-unit management expenditures

Based on the R andom samples described above, we applied linear models using the base lm() function from R v4.0.3 (R Core Team 2019), to compute the coefficients of equation 8 from the 36 data points corresponding to the species by biome combinations. We quantified I as half of the management cost between 1995 and 2008 (since it represents twice the period over which the extent of the treated area was provided). R was quantified as the condensed treated area per species and per biome, and A as the mean abundance in the sampled records. The per-unit management expenditures E_0_ of the different species in the biomes they were managed were then calculated using equation 9.

Random sampling of records and regressions was performed 1000 times for each sampling approach (R andom or weighted sampling), selecting Parea_S,B_% of the records each time. That means that between replicates, I and R are constant, but A varies; consequently, the approach generates a distribution of E_0_ values that reflects the variance of abundance between records for each species × biome combination. Normality of the residuals was assessed using Q-Q plots (Figs S1-S3).

## 3. Results

### 3.1. Relationship between total money spent, range size and local abundance, computed across all species combined (α and β coefficients)

Results are presented for the R andom allocation of management, management weighted by local abundance of SAPIA records, and management weighted by the inverse of local abundance of SAPIA records respectively, after applying the multi-species method. The median variance explained by the linear model over the 1000 replicates was 0.86 (adjusted r^2^, Figs S4-S6., slightly higher for management weighted by local abundance). α coefficients for range size were always significant (p values < 0.001). By contrast, only 0.3%, 7.6% and 5% of β coefficients for abundance were significant across replicates (p values ≤ 0.05). Coefficients were always above 1 for range size (after rescaling values), indicating an accelerating positive relationship with management cost (type III relationship, Fig. 1). 41%, 97.1% and 12.7% of the coefficients were above 0 for abundance. None of the values below 0 was significant. However, the sampling approach influenced how the coefficient values were distributed around 1. For the R andom sampling, 66% of the significant coefficients were below 1 (type II decelerating relationship; Figs 1, S4). For the sampling weighted by local abundance of SAPIA records, all significant coefficients were above 1 (type III accelerating relationship; Figs 1, S5). For the sampling weighted by the inverse of local abundance of SAPIA records, 98% of the significant coefficients were below 1 (type II decelerating relationship; Figs 1, S6).

### 3.2. Species-specific per-unit expenditures (E_0,s_)

We only report money spent per-unit for positive α and β coefficients, as negative values should be taken as artefacts from stochastic sampling. The money spent per-unit obtained after applying the multi-species method was qualitatively similar across the different approaches for allocating management effort. Since the models fitted from using the weighted sampling of the SAPIA records generated a very slightly higher explained variance, and coefficients were more consistently positive and significant than for the other two approaches, we will present these results in the following (Fig. 4, but see Figs S7 and S8 for the money spent per-unit calculated with the other two approaches to allocating management).

**Figure 4.**
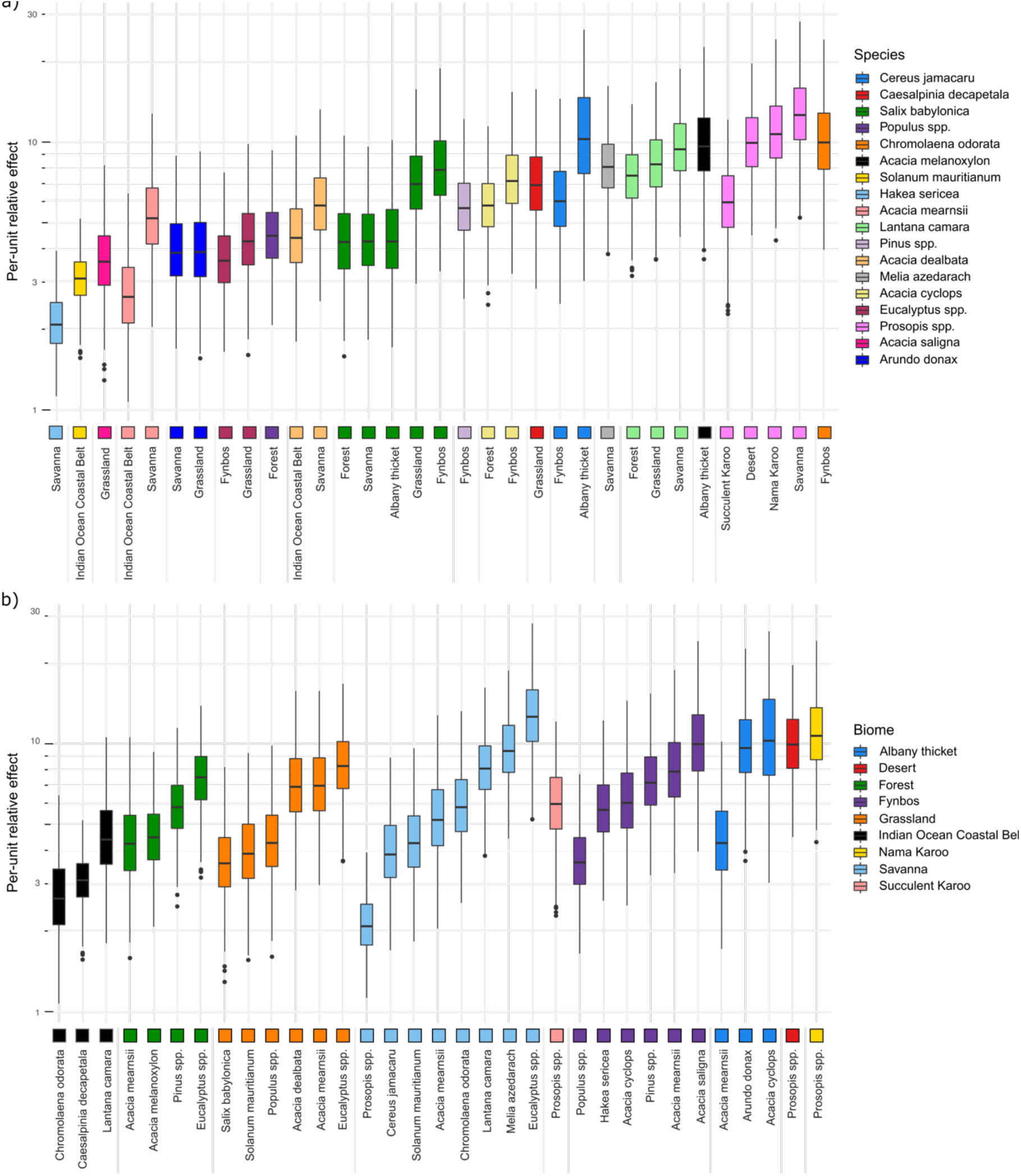
Distributions of per-unit relative effects (here money spent per-unit on management) of invasive plant species managed by the Working for Water program in South Africa between 1999 and 2008, in the different biomes of South Africa, over the 1000 replicates, calculated after sampling predominantly records with high abundance. Money spent per-unit should be interpreted in a relative rather than absolute fashion, due to the lack of absolute meaning for the abundance values. a) Species are distinguished by different colours and ordered by their median money-spent per-unit over all biomes where they were managed. b) Biomes are distinguished by different colours and ordered by their median per-unit cost over all managed species they contain.

The relative money spent per-unit on the different species (Fig. 4a) differed by five orders of magnitude between invasions where the least money was spent (*Cereus jamacaru* in Savanna) to those where the most money was spent (*Prosopis* species in Savanna). Some taxa showed differences in the money spent per-unit in different biomes, for example, the money spent per-unit on *Chromolaena odorata* was about twice as high in the Savanna than in the Indian Ocean costal belt biome. The money spent per-unit on *Acacia mearnsii* was ∼1.5 times higher in the Fynbos and Grassland biomes than in the Albany thicket, Forest and Savanna biomes. The money spent per-unit on *Acacia cyclops* was also ∼1.5 times higher in the Albany thicket biomes than in the Fynbos. The money spent per-unit on *Prosopis* species was ∼1.5 times higher in the Desert, Nama Karoo, and Savanna biomes than in the Succulent Karoo. The biomes in which money spent per-unit was higher were therefore not consistent across taxa. We observed no substantial differences of money spent per-unit across biomes overall, all taxa considered (Fig. 4b).

## 4. Discussion

The application of G-IRAE shows that a straightforward regression approach facilitates the calculation of per-unit effects of alien taxa, thereby also disentangling contributions of local abundance and range size to total impact. Management of protected areas simultaneously invaded by multiple alien species is often area-, rather than species-based (Cheney et al. 2021). However, area-based approaches can be less efficient for specific high-priority species that need to be targeted (Foxcroft et al. 2009). By computing species-specific per-unit effects, our approach can complement area-based approaches and inform management from an economic perspective. At similar abundances and range size, a species with a lower per-unit management cost E_0_ should be cheaper to manage. From a species-based management perspective, it may therefore be advantageous to manage priority species with high per-unit management costs or impacts that occur at low abundance and/or over narrow ranges, before they start spreading. In addition, disentangling abundance, range size and per-unit effects will enable the factors driving each of the three components of impact to be investigated. This will ultimately improve our ability to predict future impacts of recently introduced alien species.

### 4.1. Generalising and interpreting the calculation of per-unit effects for different types of impacts

To illustrate our approach, we focused on money spent on managing species. This specific case study in part reflected data availability: money spent is routinely documented and such data are available for a wide range of species across different regions and across different countries (Diagne et al. 2020, 2021). Note that money spent is not necessarily correlated with metrics of damage nor of ecological impact. The amount of money spent is indeed often determined independently of ecological impacts, as it is affected by feasibility, return on investment, and, in the case of Working for Water, where it is most important to create jobs in order to alleviate poverty (van Wilgen and Wannenburgh 2016). The approach can nonetheless be extended to other measures of expenditure (e.g. work time spent for alien species management) or impact types, e.g. related to damage costs, human welfare, health or ecological impacts. It is important to acknowledge that alien species can impact multiple sectors of society, each with their own way of valuing goods and services, and different types of impacts will likely be more consistently documented than others (Diagne et al. 2020, 2021).

Environmental or social impacts will be more challenging to integrate in the formula, due to the complexity of these concepts and the paucity of data for assessing impacts at different spatial scales in these domains. Multiple measures of impact may be used, at the population, community or ecosystem level, and their suitability can vary depending on the life forms of the alien species (see e.g. the measures used in multiple analyses reported by Norbury et al., 2015). Global classification of impact into ordered categories, such as the EICAT scheme (Blackburn et al. 2014), endorsed by the IUCN, offer avenues to compare species globally, but the use of ordered categories in linear and generalised linear models is not straightforward (Guisan and Harrell 2000). This analytical framework therefore represents a basis from which to develop more specific statistical tools to integrate different types of data. We also hope it will incentivise systematic collection and harmonisation of impact data with data on range size and local abundance. Doing so will allow to assess and rank impacts and costs of biological invasions, and further feed decision making initiatives (many forms of analysis are routinely used as inputs into decision making; McGeoch et al. 2016). We discuss below different aspects related to range size and abundance that are important to consider when collecting data.

The time scale of impacts varies across species; some impacts occur very quickly whereas others can take decades or longer to manifest, leading to a slow accumulation of “invasion impact debt” (sensu Rouget et al., 2016). That means that E can be considered as an increasing function of time t since introduction. To account for such time lags, time since introduction could therefore be incorporated into equation 3 as follows:

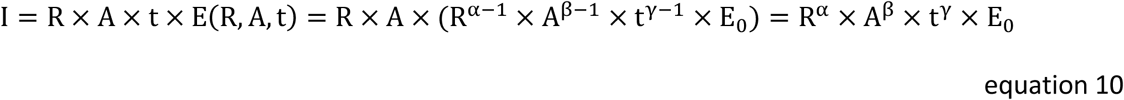

where γ determines how E increases in time.

Abundance is not a straightforward variable to monitor in a comparable fashion for multiple species, especially for plants (Catford et al. 2012). In absolute values, abundance can be measured as numbers of individuals, cover area, or unit of biomass. Number of individuals is a suitable metrics for large animals, but biomass may be more appropriate for small animals such as insects. In the case of plants, number of individuals may be more intuitive for large trees, whereas cover area is more appropriate for herbaceous plants (Wilson et al. 2014). Relative abundance (i.e. compared to the total abundance over all species present at a site) may also be appropriate, especially in relationship with ecological impacts on other species. For example, Wilson et al. (2018) propose the following definition for each class: “Invasive plants cover x - y% of the area covered by plants, or invasive species make up x - y% of the biomass of the area; populations of invasive animals make up x - y% of all individual animals at the site.” Such categories (similar to those we used here) offer a convenient way to combine these different quantities for multiple taxonomic groups, although they imply that the per-unit effects computed with our method should also be interpreted in a relative rather than absolute fashion. This also implies that for results to be comparable across different regions, the same classification scheme should be used.

Although more straightforward than abundance, range size in Parker et al.’s (1999) formula (equation 1) may also be computed in different ways. Two main measures are typically used in conservation science across taxonomic groups: the extent of occurrence (EoO) and the area of occupancy (AoO) (IUCN 2001) (but see Hui et al., 2011 for more complex alternatives). The EoO is the minimum continuous area encompassing all sites of occurrences of a taxon (for example represented by a convex hull). However, the EoO may not be appropriate when a species is introduced to a country or region at multiple locations that may be far apart leaving large areas with no individuals (and therefore no impact) in between. The AoO is the area effectively occupied by a taxon, and corresponds to the condensed area used by van Wilgen et al. (2012) and in our analyses. In practice, accuracy of the AoO depends on the spatial grain at which it is estimated. The choice of a measure for computing the per-unit effect of an alien species will be case-dependent, based on the type of impact and the context. For example, in the case of management costs, if the same team is to control an alien species population, it may be costlier to control a patchy population spread over a larger area than a population occurring over a small area, due to the time necessary to move from one invaded location to another. In that case, the EoO provides additional information not captured by the AoO.

### 4.2. Predicting alien species impacts

Disentangling the three components of impact and characterising alien species by combinations of the three coefficients (*α, β*, E_0_) is an essential first step towards identifying which species and environmental attributes determine the values of the three coefficients. Doing so will enable prediction of the potential impacts of newly introduced species or species on watch lists, or their future management costs. For ecological impacts, the predictors linked to per-unit effect can be related to the constituents of the environment, i.e. attributes of the invader, attributes of the resident species, resources levels, and abiotic conditions (Thiele et al. 2010); they can also be attributes of the interactions between these constituents, i.e. propagule and colonisation pressure, functional distinctiveness, environmental tolerance, interactions with resident species, disturbance, and environmental heterogeneity (Ricciardi et al. 2013). For per-unit management costs, the attributes of the species are important (e.g. a large tree may be more expensive to remove than a shrub; resprouting species are usually more difficult to control than those that regenerate only from seeds, etc., although we could not detect such effects in our analyses). The treatment that is applied (e.g. mechanical removal versus herbicide application), the terrain and accessibility of the location where populations occur, etc., should also be considered (Panetta 2009).

Different combinations of predicted range size, local abundance, α and β coefficients, and E_0_ values can then suggest different levels of concern for the corresponding species. For example, for a given maximum impact I, a species with a type III relationship (*α* or *β* above 1) will only be of concern if range size is predicted to be large or local abundance high, as the total impact would remain low at low range size and abundance. In contrast, for species with a type II relationship (*α* or *β* below 1) and the same maximum impact I at the same R and A values (and therefore a higher per-unit effect), the predicted range size or abundance has little importance (since impact mostly varies at low R or A values in this case), and concern will be mostly determined by the per-unit effects of species and the resulting maximum impact they can have.

### 4.3. Limitations of the approach and solutions

Here, management expenditures were reported at the scale of biomes, and some species were managed in one or two biomes. As a result, only the multi-species method could be applied. However, effects can be assessed at other spatial scales. Spatial units of assessment can encompass lakes (Latzka et al. 2016), patches of several thousands of hectares (Holland et al. 2013), etc. Fine-scale assessments of impacts at multiple locations, congruently with local abundance and area of the location, recorded for multiple species, would allow for a broader application of the species-specific method. Different types of impacts would likely be measurable at different spatial scales, and results should be interpreted in light of the scale.

The multi-species method illustrated in this paper generates a fixed per-unit money spent E_0_ for a species in a given environment, that is independent of its abundance and range size. The approach assumes that the relationship between money spent or impact and range size or local abundance is the same for all species in the different biomes. This is a necessary simplification, as comprehensive data on species range, local abundance and money spent or impact in multiple locations required for the species-specific method are missing for most species. The high variance explained by the model nonetheless suggests that this is a valid approximation in our case study on money spent per species in different biomes. As a result, E_0_ represents the per-unit money spent of a species in the absence of conspecifics (i.e. when it is only present in a unit of range size by a unit of abundance), therefore allowing to compare alien species irrespectively of their abundance or range size.

If data are not available at the species level, but different species are expected to be characterised by different relationships between impact, abundance and range size, a compromise would be to apply equations 6 – 8 to different groups of species, for which the relationship between per-unit effect, range size and abundance is assumed to differ between groups, but to be the same within groups. Grouping species according to these relationships will likely be a challenge, and should be guided by expert knowledge on the biology of the species (e.g. invasion syndromes; Novoa et al., 2020; Perkins & Nowak, 2013).

### 4.4. Money spent managing invasive plants in South Africa

We found that species identity played an important role in explaining the money spent per-unit, although we could not identify species attributes that may explain these differences. Differences in per-unit money spent between biomes were also not consistent across the taxa present in multiple biomes. This was probably due to the spatial grain of biomes being too large, and each therefore encompasses a wide variety of terrains and conditions which influence money spent per-unit.

Nonetheless, the clear differences between the money spent per-unit on the same taxa in different biomes suggests that the differences in protocols and methods used for their management in different biomes warrant further investigation. This may be due to genuine differences in the ease of management of particular species between biomes, reflect fundamental differences in the relationships between abundance and range size in different environments, or emerge from differences in the optimisation of management of multiple alien species simultaneously in the biomes.

Finally, it is important to note that we did not have data on the outcome of the management, i.e. whether the money spent on control was effective in reducing the invasive plant populations. Our analysis looked only at broad-scale inferences of occupied areas before and after the Working for Water assessment, which were performed using different methods. As such, the comparisons as described above are in terms of money spent per-unit rather than costs to manage (i.e. how much it would cost to keep an invasion below a threshold). Explicit assessments of management efficacy are rare in South Africa (Zengeya & Wilson, 2020, although see e.g. Kraaij et al., 2017).

## 5. Conclusions

We have proposed G-IRAE, a simple approach for calculating the per-unit effects of alien species, including ecological impacts, damage costs, or management expenditures. This approach provides the means to disentangle the per-unit effect from those of range size and local abundance to total impact for a wide range of taxonomic groups. We illustrated it for plant data using money spent on management, but other taxonomic groups and types of impact can be considered, using appropriate metrics for range size and local abundance. Two methods, the species-specific and the multi-species methods, can be used depending on the available data for each species. G-IRAE provides a tool for exploring how characteristics of the alien species and recipient environments, and their interactions, influence different components of impact. Such insights are crucial for designing and prioritising appropriate management actions, at both the regional and national scales, and moving towards objective frameworks for anticipating and responding to the impacts of alien species.

## Acknowledgements

We thank Franck Courchamp and Christophe Diagne for useful comments on a preliminary version of this manuscript. MAM acknowledges support from the Australian Research Council (DP200101680). This research was funded through the 2017-2018 Belmont Forum and BiodivERsA joint call for research proposals, under the BiodivScen ERA-Net COFUND programme, and with the funding organisation Austrian Science Foundation FWF (grant I 4011-B32) (to FE and BL). DMR acknowledges support from the DSI-NRF Centre of Excellence for Invasion Biology, the Oppenheimer Memorial Trust (grant 18576/03) and the Millennium Trust. JRUW thanks the South African Department of Forestry, Fisheries, and the Environment (DFFE) for funding, noting that this publication does not necessarily represent the views or opinions of DFFE or its employees. JAC acknowledges funding from the European Research Council (ERC) under the European Union’s Horizon 2020 research and innovation programme (grant agreement No. [101002987]).

## Data accessibility

Data from van Wilgen et al. (2012) is directly accessible from the original article. The SAPIA database can be accessed by contacting the South Africa National Biodiversity Institute – SANBI (http://biodiversityadvisor.sanbi.org/contact/).

## Author contributions

GL and MAM conceived the original concept. GL designed and performed the analyses and drafted the manuscript. All authors contributed to the refinement of the analyses, to the interpretation of the results, and to the multiple rewriting rounds of the manuscript.

## Notes

### Competing Interest Statement

The authors have declared no competing interest.

## References

Bellard C, Cassey P, Blackburn TM (2016) Alien species as a driver of recent extinctions. Biol Lett 12:20150623. https://doi.org/10.1098/rsbl.2015.0623

Blackburn TM, Essl F, Evans T, et al (2014) A unified classification of alien species based on the magnitude of their environmental impacts. PLoS Biol 12:e1001850. https://doi.org/10.1371/journal.pbio.1001850

Bradley BA, Laginhas BB, Whitlock R, et al (2019) Disentangling the abundance–impact relationship for invasive species. Proc Natl Acad Sci 116:9919–9924

Buckley YM, Catford J (2016) Does the biogeographic origin of species matter? Ecological effects of native and non-native species and the use of origin to guide management. J Ecol 104:4–17. https://doi.org/10.1111/1365-2745.12501

Catford JA, Vesk PA, Richardson DM, Pyšek P (2012) Quantifying levels of biological invasion: towards the objective classification of invaded and invasible ecosystems. Glob Chang Biol 18:44–62. https://doi.org/10.1111/j.1365-2486.2011.02549.x

Cheney C, van Wilgen NJ, Esler KJ, et al (2021) Quantifying range structure to inform management in invaded landscapes. J Appl Ecol 58:338–349. https://doi.org/10.1111/1365-2664.13765

Cuthbert RN, Dickey JWE, Coughlan NE, et al (2019) The Functional Response Ratio (FRR): advancing comparative metrics for predicting the ecological impacts of invasive alien species. Biol Invasions 21:2543–2547. https://doi.org/10.1007/s10530-019-02002-z

Diagne C, Leroy B, Gozlan RE, et al (2020) InvaCost, a public database of the economic costs of biological invasions worldwide. Sci Data 7:1–12. https://doi.org/10.1038/s41597-020-00586-zgas

Diagne C, Leroy B, Vaissière A-C, et al (2021) High and rising economic costs of biological invasions worldwide. Nature. https://doi.org/10.1038/s41586-021-03405-6

Foxcroft LC, Richardson DM, Rouget M, MacFadyen S (2009) Patterns of alien plant distribution at multiple spatial scales in a large national park: implications for ecology, management and monitoring. Divers Distrib 15:367–378. https://doi.org/10.1111/j.1472-4642.2008.00544.x

Guisan A, Harrell FE (2000) Ordinal response regression models in ecology. J Veg Sci 11:617–626. https://doi.org/10.2307/3236568

Henderson L (2007) Invasive, naturalized and casual alien plants in southern Africa: a summary based on the Southern African Plant Invaders Atlas (SAPIA). Bothalia 37:215–248. https://doi.org/10.4102/abc.v37i2.322

Henderson L, Wilson JRU (2017) Changes in the composition and distribution of alien plants in South Africa: An update from the Southern African Plant Invaders Atlas. Bothalia-African Biodivers Conserv 47:1–26. http://dx.doi.org/10.4102/abc.v47i2.2172

Holland EP, Pech RP, Ruscoe WA, et al (2013) Thresholds in plant–herbivore interactions: predicting plant mortality due to herbivore browse damage. Oecologia 172:751–766. https://doi.org/10.1007/s00442-012-2523-5

Hui C, Richardson DM, Robertson MP, et al (2011) Macroecology meets invasion ecology: linking the native distributions of Australian acacias to invasiveness. Divers Distrib 17:872–883. https://doi.org/10.1111/j.1472-4642.2011.00804.x

IPBES (2019) Global assessment report on biodiversity and ecosystem services. IPBES Secretariat, Bonn, Germany

IUCN (2001) IUCN Red List categories and criteria. IUCN, Gland, Switzerland and Cambridge, UK

Jeschke JM, Bacher S, Blackburn TM, et al (2014) Defining the impact of non-native species. Conserv Biol 28:1188–1194

Kotzé JDF, Beukes BH, Van den Berg EC, Newby TS (2010) National invasive alien plant survey. Report Number: GW/A/2010/21. Agricultural Research Council: Institute for Soil, Climate and Water, Pretoria

Kraaij T, Baard JA, Rikhotso DR, et al (2017) Assessing the effectiveness of invasive alien plant management in a large fynbos protected area. Bothalia - African Biodivers. Conserv. 47:1–11

Kumschick S, Bacher S, Bertolino S, et al (2020) Appropriate uses of EICAT protocol, data and classifications. NeoBiota 62:193. https://doi.org/10.3897/neobiota.62.51574

Latombe G, Essl F, McGeoch MA (2020) The effect of cross-boundary management on the trajectory to commonness in biological invasions. NeoBiota 62:241–267

Latzka AW, Hansen GJA, Kornis M, Vander Zanden MJ (2016) Spatial heterogeneity in invasive species impacts at the landscape scale. Ecosphere 7:e01311. https://doi.org/10.1002/ecs2.1311

Le Maitre DC, Versfeld DB, Chapman RA (2000) Impact of invading alien plants on surface water resources in South Africa: A preliminary assessment. Water South Africa 26:397–408

Maxwell SL, Fuller RA, Brooks TM, Watson JEM (2016) Biodiversity: The ravages of guns, nets and bulldozers. Nat News 536:143. https://doi.org/10.1038/536143a

McGeoch MA, Genovesi P, Bellingham PJ, et al (2016) Prioritizing species, pathways, and sites to achieve conservation targets for biological invasion. Biol Invasions 18:299–314. https://doi.org/10.1007/s10530-015-1013-1

McGeoch MA, Latombe G (2016) Characterizing common and range expanding species. J Biogeogr 43:217–228. https://doi.org/10.1111/jbi.12642r

Norbury GL, Pech RP, Byrom AE, Innes J (2015) Density-impact functions for terrestrial vertebrate pests and indigenous biota: Guidelines for conservation managers. Biol Conserv 191:409–420. https://doi.org/10.1016/j.biocon.2015.07.031

Novoa A, Richardson DM, Pyšek P, et al (2020) Invasion syndromes: A systematic approach for predicting biological invasions and facilitating effective management. Biol Invasions 1–20

Panetta FD (2009) Weed Eradication—An Economic Perspective. Invasive Plant Sci Manag 2:360– 368. https://doi.org/10.1614/IPSM-09-003.1

Parker IM, Simberloff D, Lonsdale WM, et al (1999) Impact: toward a framework for understanding the ecological effects of invaders. Biol Invasions 1:3–19. https://doi.org/10.1023/A:1010034312781

Perkins LB, Nowak RS (2013) Invasion syndromes: hypotheses on relationships among invasive species attributes and characteristics of invaded sites. J Arid Land 5:275–283

Pyšek P, Richardson DM, Jarošík V (2006) Who cites who in the invasion zoo: insights from an analysis of the most highly cited papers in invasion ecology. Preslia 78:437–468

R Core Team (2019) R: A language and environment for statistical computing

Ricciardi A, Hoopes MF, Marchetti MP, Lockwood JL (2013) Progress toward understanding the ecological impacts of nonnative species. Ecol Monogr 83:263–282. https://doi.org/10.1890/13-0183.1

Rouget M, Robertson MP, Wilson JRU, et al (2016) Invasion debt–Quantifying future biological invasions. Divers Distrib 22:445–456

Roura-Pascual N, Krug RM, Richardson DM, Hui C (2010) Spatially-explicit sensitivity analysis for conservation management: exploring the influence of decisions in invasive alien plant management. Divers Distrib 16:426–438. https://doi.org/10.1111/j.1472-4642.2010.00659.x

Roura-Pascual N, Richardson DM, Krug RM, et al (2009) Ecology and management of alien plant invasions in South African fynbos: Accommodating key complexities in objective decision making. Biol Conserv 142:1595–1604. https://doi.org/10.1016/j.biocon.2009.02.029

Seidl R, Klonner G, Rammer W, et al (2018) Invasive alien pests threaten the carbon stored in Europe’s forests. Nat Commun 9:1–10. https://doi.org/10.1038/s41467-018-04096-w

Shackleton RT, Le Maitre DC, van Wilgen BW, Richardson DM (2017) Towards a national strategy to optimise the management of a widespread invasive tree (Prosopis species; mesquite) in South Africa. Ecosyst Serv 27:242–252. https://doi.org/10.1016/j.ecoser.2016.11.022

South African National BiodiversityInstitute (2006) Vegetation Map of South Africa, Lesotho and Swaziland 2006 [vector geospatial dataset]. Available from the Biodiversity GIS website. Accessed 5 Jan 2021

Strayer DL (2020) Non-native species have multiple abundance–impact curves. Ecol Evol 10:6833– 6843. https://doi.org/10.1002/ece3.6364

Thiele J, Kollmann J, Markussen B, Otte A (2010) Impact assessment revisited: improving the theoretical basis for management of invasive alien species. Biol Invasions 12:2025–2035. https://doi.org/10.1007/s10530-009-9605-2

Van den Berg EC, Kotze I, Beukes H (2013) Detection, quantification and monitoring of Prosopis in the Northern Cape Province of South Africa using remote sensing and GIS. South African J Geomatics 2:68–81

Van der Colff D, Kumschick S, Foden W, Wilson JRU (2021) Comparing the IUCN’s EICAT and Red List to improve assessments of the impact of biological invasions. NeoBiota 62:509–523

van Wilgen BW, Forsyth GG, Le Maitre DC, et al (2012) An assessment of the effectiveness of a large, national-scale invasive alien plant control strategy in South Africa. Biol Conserv 148:28–38. https://doi.org/10.1016/j.biocon.2011.12.035

van Wilgen BW, Wannenburgh A (2016) Co-facilitating invasive species control, water conservation and poverty relief: achievements and challenges in South Africa’s Working for Water programme. Curr Opin Environ Sustain 19:7–17. https://doi.org/10.1016/j.cosust.2015.08.012

Vander Zanden MJ, Hansen GJA, Latzka AW (2017) A framework for evaluating heterogeneity and landscape-level impacts of non-native aquatic species. Ecosystems 20:477–491. https://doi.org/10.1007/s10021-016-0102-z

Wilson JRU, Caplat P, Dickie IA, et al (2014) A standardized set of metrics to assess and monitor tree invasions. Biol Invasions 16:535–551

Wilson JRU, Datta A, Hirsch H, et al (2020) Is invasion science moving towards agreed standards? The influence of selected frameworks. NeoBiota 62:569–590

Wilson JRU, Faulkner KT, Rahlao SJ, et al (2018) Indicators for monitoring biological invasions at a national level. J Appl Ecol doi: 10.1111/1365-2664.13251

Yokomizo H, Possingham HP, Thomas MB, Buckley YM (2009) Managing the impact of invasive species: the value of knowing the density–impact curve. Ecol Appl 19:376–386. https://doi.org/10.1890/08-0442.1

Zengeya TA, Wilson JR (eds) (2020) The status of biological invasions and their management in South Africa in 2019

