## Supplementary figures for "G-IRAE: a Generalised approach for linking the total Impact of invasion to species’ Range, Abundance and per-unit Effects"

This document contains additional figures presenting results from analyses.

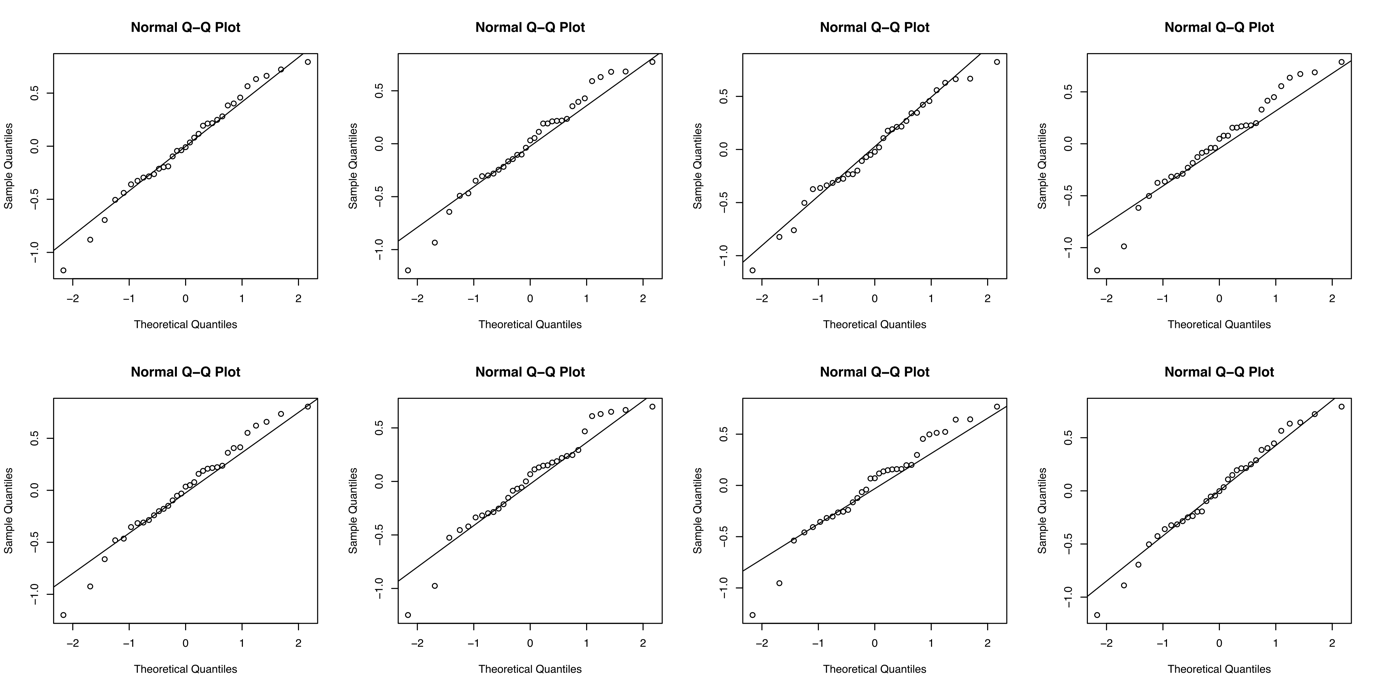

**Figure S1.** Q-Q plots of model residuals for eight random replicates out of 1000, calculated after randomly sampling SAPIA records.

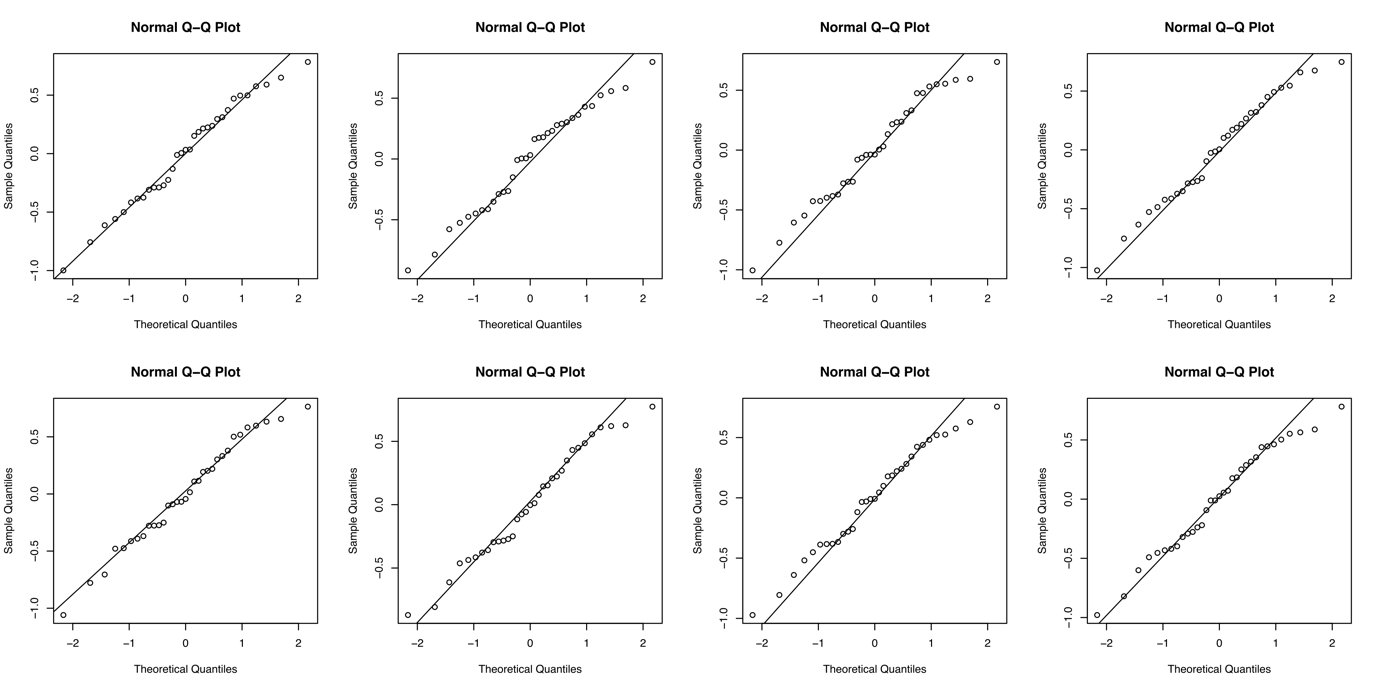

**Figure S2.** Q-Q plots of model residuals for eight random replicates out of 1000, calculated after sampling predominantly SAPIA records with high abundance.

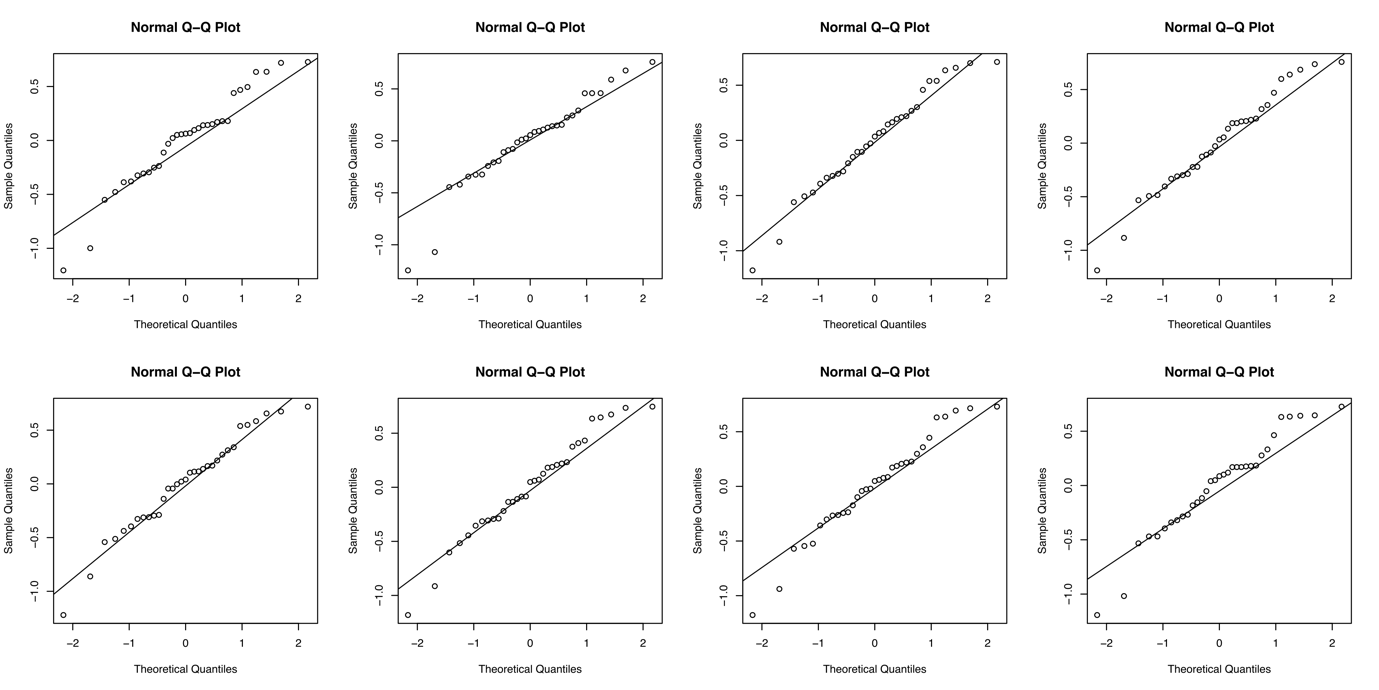

**Figure S3.** Q-Q plots of model residuals for eight random replicates out of 1000, calculated after sampling predominantly SAPIA records with low abundance.

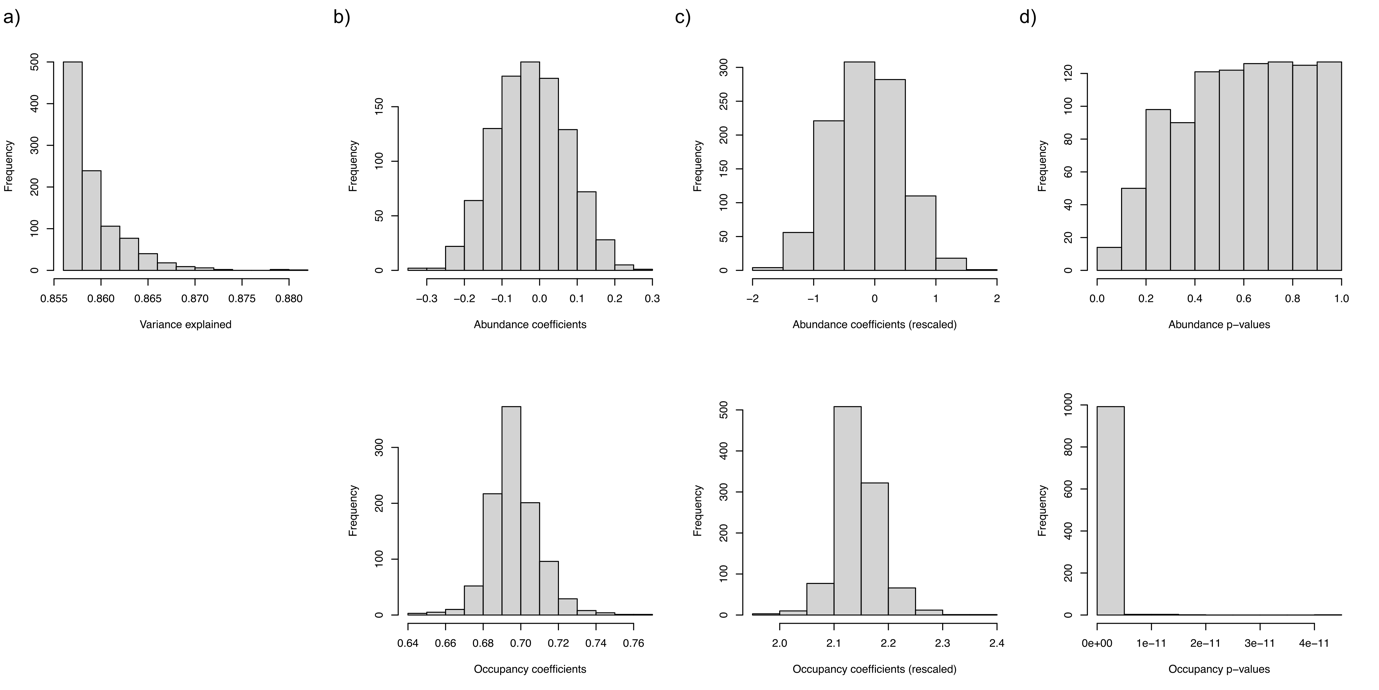

**Figure S4.** Model results calculated after randomly sampling SAPIA records. a) Distribution of the variance explained by the models for the 1000 replicates. b) Distributions of the α and β coefficients for abundance and occupancy. c) Distributions of the α and β coefficients for abundance and occupancy after rescaling values between [0 – 1]. d) Distributions of the p-values of the α and β coefficients.

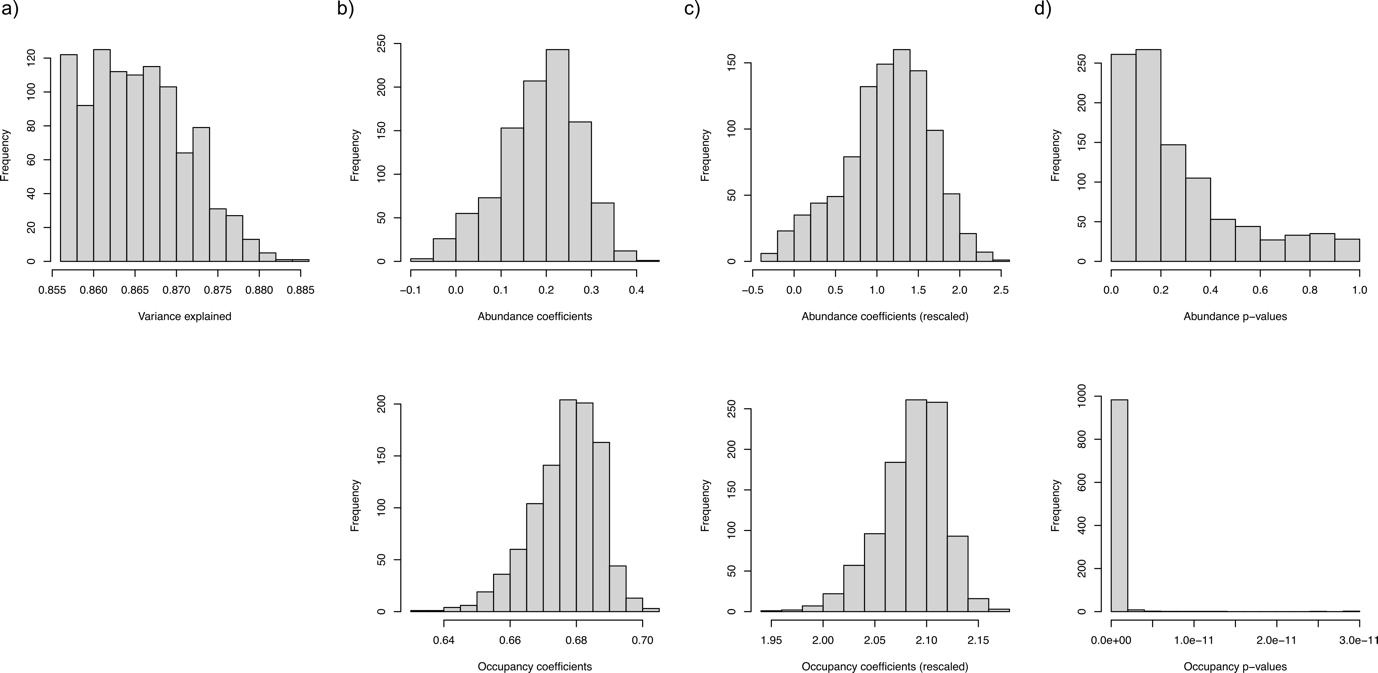

**Figure S5.** Model results calculated after sampling predominantly SAPIA records with high abundance. a) Distribution of the variance explained by the models for the 1000 replicates. b) Distributions of the α and β coefficients for abundance and occupancy. c) Distributions of the α and β coefficients for abundance and occupancy after rescaling values between [0 – 1]. d) Distributions of the p-values of the α and β coefficients.

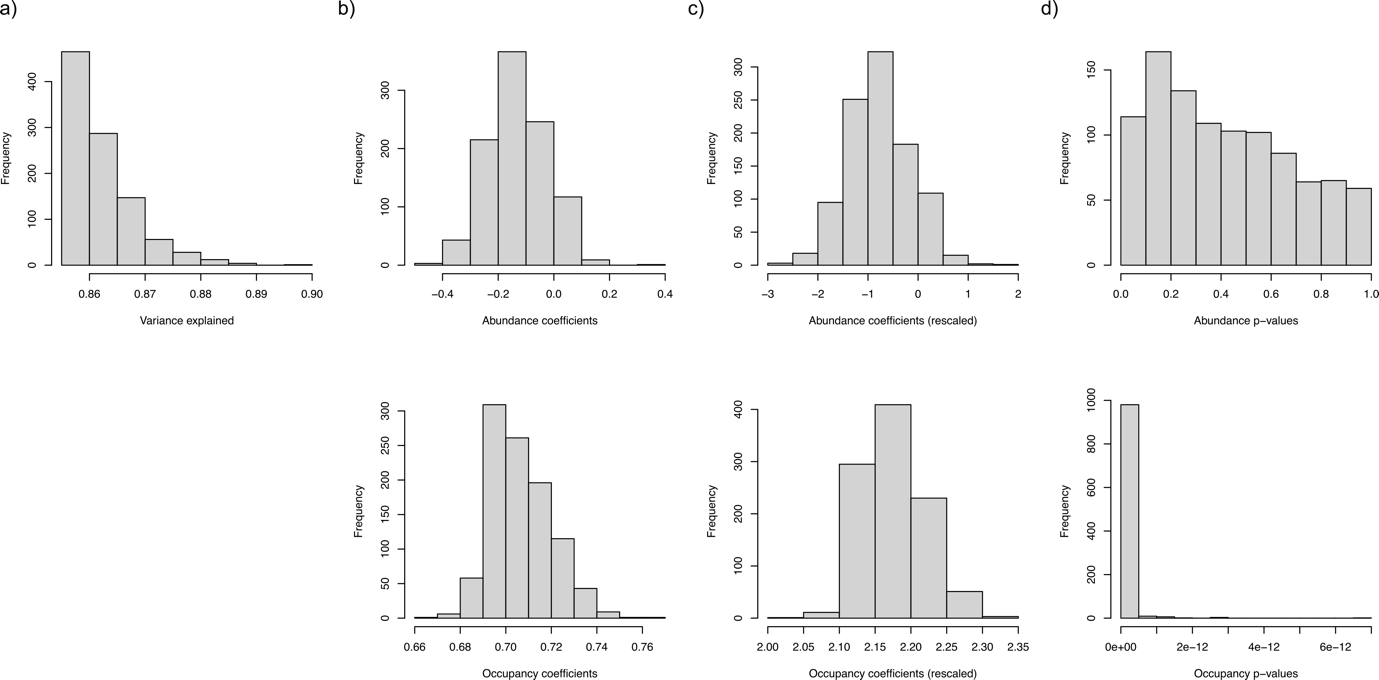

**Figure S6.** Model calculated after sampling predominantly SAPIA records with low abundance. a) Distribution of the variance explained by the models for the 1000 replicates. b) Distributions of the α and β coefficients for abundance and occupancy. c) Distributions of the α and β coefficients for abundance and occupancy after rescaling values between [0 – 1]. d) Distributions of the p-values of the α and β coefficients.

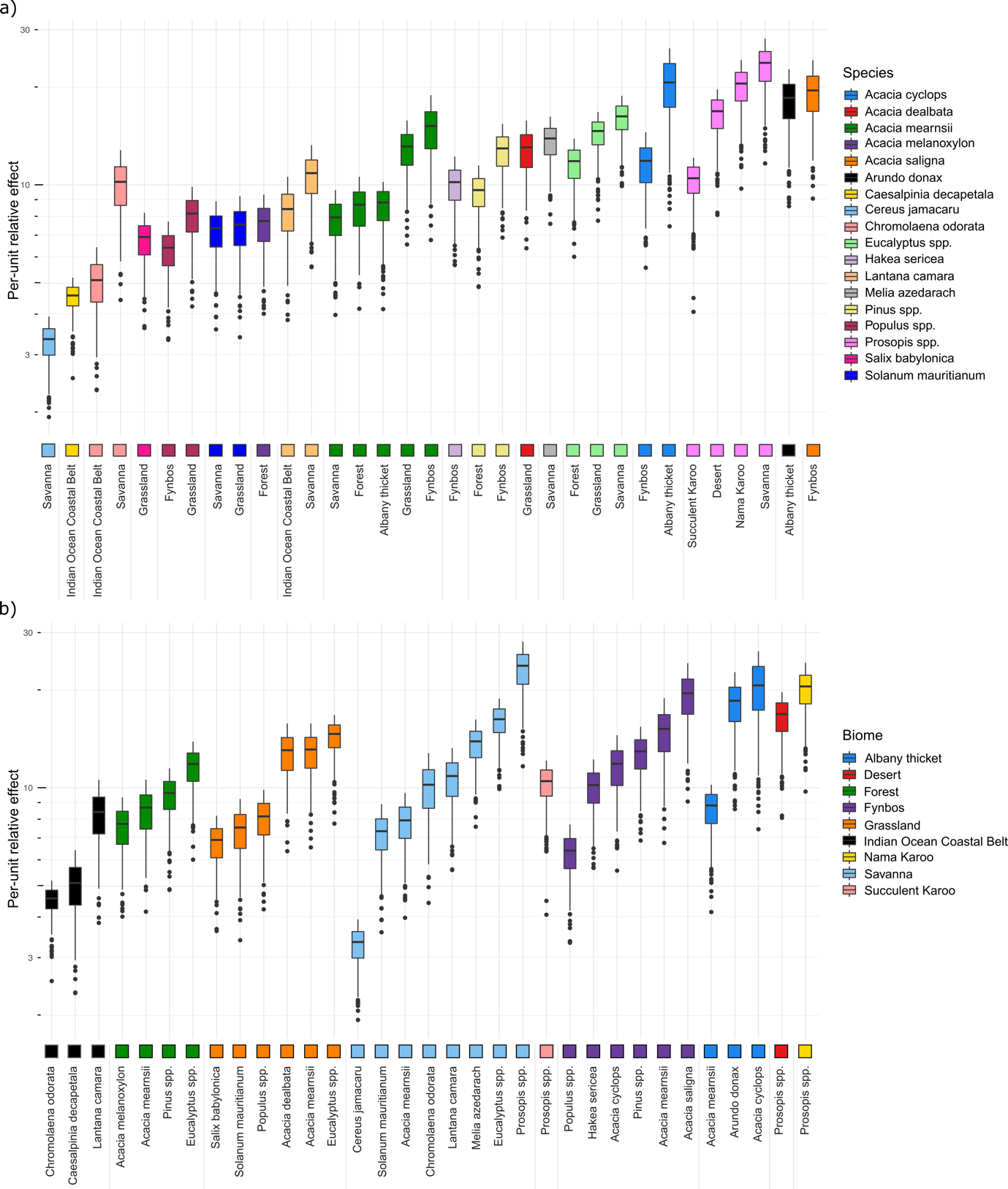

**Figure S7.** Distributions of per-unit relative effects (here per-unit relative management costs, i.e. money spent per-unit on management) of alien species managed by the Working for Water program in South Africa between 1999 and 2008, in the different biomes of South Africa, over the 1000 replicates, calculated after randomly sampling records. Per-unit cost values should be interpreted in a relative rather than absolute fashion, due to the lack of absolute meaning for the abundance values. a) Species are distinguished by different colours and ordered by their median per-unit cost over all biomes where they were managed. b) Biomes are distinguished by different colours and ordered by their median per-unit cost over all managed species they contain.

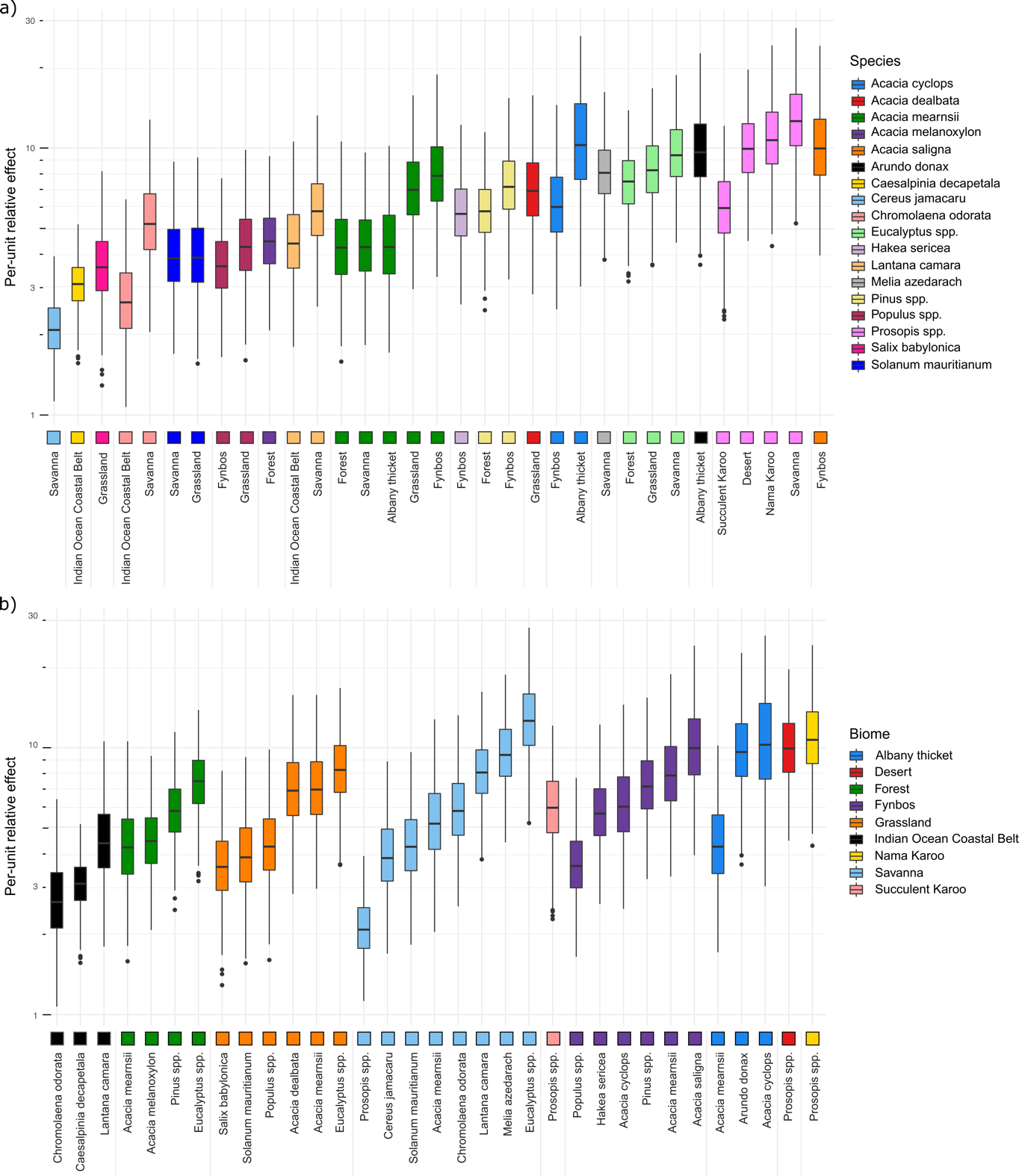

**Figure S8.** Distributions of per-unit relative effects (here per-unit relative management costs, i.e. money spent per-unit on management) of alien species managed by the Working for Water program in South Africa between 1999 and 2008, in the different biomes of South Africa, over the 1000 replicates, calculated after sampling predominantly records with low abundance. Per-unit cost values should be interpreted in a relative rather than absolute fashion, due to the lack of absolute meaning for the abundance values. a) Species are distinguished by different colours and ordered by their median per-unit cost over all biomes where they were managed. b) Biomes are distinguished by different colours and ordered by their median per-unit cost over all managed species they contain.
